# Identification of leukemia stem cell subsets with distinct transcriptional, epigenetic and functional properties

**DOI:** 10.1101/2024.02.09.579319

**Authors:** Héléna Boutzen, Alex Murison, Jean C. Y. Wang, Christopher Arlidge, Mathieu Lupien, Kerstin B. Kaufmann, John E. Dick

## Abstract

The leukemia stem cell (LSC) compartment is a complex reservoir fueling disease progression in acute myeloid leukemia (AML). The existence of heterogeneity within this compartment is well documented but prior studies have focused on genetic heterogeneity without being able to address functional heterogeneity. Understanding this heterogeneity is critical for the informed design of therapies targeting LSC, but has been hampered by LSC scarcity and the lack of reliable cell surface markers for viable LSC isolation. To overcome these challenges, we turned to the patient-derived OCI-AML22 cell model. This model includes functionally, transcriptionally and epigenetically characterized LSC broadly representative of LSC found in primary AML samples. Focusing on the pool of LSC, we used an integrated approach combining xenograft assays with single-cell analysis to identify two LSC subtypes with distinct transcriptional, epigenetic and functional properties. These LSC subtypes differed in depth of quiescence, differentiation potential and repopulation capacity and could be isolated based on CD112 expression. A majority of AML patient samples had transcriptional signatures reflective of either LSC subtype, and some even showed coexistence within an individual sample. This work provides a framework for further investigation of the LSC compartment in AML.

## Introduction

LSC have been tightly linked to disease progression and poor prognosis in AML, and the development of more effective therapies to eradicate LSC is an important unmet need^1–4^. However, progress has been hampered by inter- and intra-patient heterogeneity within the LSC compartment together with a limited ability to resolve intricacies within the pool of LSC^5^. Studies of the heterogeneity within the normal hematopoietic stem cell (HSC) compartment in mice and humans^6^ ^7–9^ provide a framework for investigating heterogeneity within the LSC pool. One of the key tools that facilitated a deeper understanding of the HSC compartment at the functional and molecular level was the identification of cell surface markers enabling purification of human HSC at nearly single cell resolution^10^. For example, immunophenotypic mapping of the HSC compartment coupled with single cell functional repopulation assays resulted in the identification of three HSC compartments: long-term (LT-), intermediate and short-term HSC with the LT-HSC subset being the most primitive ^6^ ^7–9^. Recently, we showed that LT-HSC can be further fractionated based on expression of the cell surface marker CD112 into subsets that are molecularly, transcriptionally and functionally distinct^11^. In xenograft repopulation assays, CD112-Low HSC exhibit latency, with slower engraftment kinetics compared to CD112-High HSC^12^. However, CD112-Low HSC outcompete CD112-High HSC in long-term upon serial transplantation^11^.

In contrast to the normal HSC compartment, there are few tools currently available to study the LSC compartment. The existence of heterogeneity within this compartment is well documented in leukemia^13–19^, but prior studies have focused on genetic mutations without being able to address functional heterogeneity. More than two decades ago, we used clonal tracking approaches to show that the LSC pool comprises multiple LSC subtypes with distinct functional capacities, with some LSC more latent and slowly repopulating in xenograft assays and others exhibiting faster repopulation kinetics^20^. Given that the existence of functional heterogeneity among LSCs is likely associated with differences in therapeutic vulnerabilities, a better understanding of the basis for the observed heterogeneity is crucial for the rational design of effective therapies aimed at eradicating the entire pool of LSC^21^. However, progress has been hampered by the lack of tools for prospective isolation of distinct LSC subtypes. Although several LSC surface markers have been described^22–25^, none reliably capture functional LSC across a broad spectrum of AML patients.

Despite this heterogeneity, transcriptomic and epigenetic signatures do provide evidence that there are core properties of LSC that are broadly applicable across the full spectrum of AML^1,2,26–28^. For example, the LSC17 stemness score is highly predictive of outcome across thousands of pediatric and adult AML samples, irrespective of molecular and cytogenetic subtype^2,29–31^. In addition, single cell RNA-seq analysis uncovered signatures of stem, progenitor and mature AML cells that enabled the development of a hierarchy classification system that predicts drug sensitivity in AML patients^32,33^. These findings suggest that core stemness properties of LSC that drive clinical outcomes and treatment sensitivity are shared across AML patients, and support studies to understand whether the basis for the observed functional heterogeneity within the LSC compartment may also be shared. The gold standard for studies of functional properties of LSC has been AML patient-derived xenograft (PDX) assays, as most AML cell lines do not exhibit a LSC-driven hierarchy that captures patient biology^34^. However, PDX assays are laborious, difficult to standardize and cannot be tailored for high-throughput studies. Recently, we developed a patient-derived AML model (OCI-AML22) that retains biological properties with broad clinical relevance^34^. The OCI-AML22 LSC population exhibits hallmark stemness properties that are highly prognostic across diverse independent cohorts of AML patient samples^1,2,29–32,35–40^. This clinically-relevant model provides a powerful tool for mechanistic and functional studies to investigate LSC biology in the correct cellular context.

Here, we used an integrated approach combining functional assays with single-cell multiome analysis in the OCI-AML22 model to identify two distinct LSC subtypes that could be prospectively isolated. These LSC subtypes possess distinct properties such as engraftment kinetics and depth of quiescence, that are retained through serial transplantation. The majority of AML patient samples possessed either of these LSC subtypes, albeit in varying proportions attesting to the broad biological relevance of our findings.

## Results

### Single cell multiome analysis captures heterogeneity within the OCI-AML22 CD34+CD38-cell fraction

To characterize in depth the pool of LSC in OCI-AML22, we performed single cell (sc) multiome analysis (scATAC-Seq/scRNA-Seq) on 7,160 cells isolated from the LSC-enriched CD34+CD38-fraction (*in vivo* LSC frequency is 1 in 200)^34^ (Figure 1A). The majority of these cells were enriched for stem cell programs when mapped onto a single cell transcriptomic map of stem and progenitor populations from healthy human bone marrow (Supplementary Figure 1A)^41^. To characterize these cells, we generated a Uniform Manifold Approximation and Projection (UMAP) based on both RNA-Seq and ATAC-Seq data and labelled cells based on their colocalization with normal hematopoietic populations when mapped onto the bone marrow reference map^41^. Interestingly, cells labelled as HSC-MPP (multipotent progenitor) - the population most enriched for self-renewing cells in the normal hematopoietic hierarchy - mapped to two distinct regions of the UMAP representation (Figure 1B). Other immature populations similarly mapped to these 2 regions (Supplementary Figure 2). Cells labelled as more committed progenitor-like, such as cycling progenitors or early GMP, mapped in between the regions occupied by HSC/MPP-like cells (Supplementary Figure 2). We also evaluated the epigenetic states of individual cells within this LSC pool using chromatin accessibility signatures from highly purified cord blood populations as a reference^42^. Using chromVar to calculate the per-cell enrichment for each signature, we identified cells highly enriched for two stem cell signatures previously identified^42^ (LT/HSPC and ACT/HSPC signatures; Figure 1 C-D), as well as cells enriched for signatures of more committed populations such as monocytes or granulocytes^42^ (Supplementary Figure 3). This analysis revealed the existence of two epigenetically distinct stem-like cell clusters localizing to the two regions occupied by transcriptionally characterized stem-like cells (Figure 1 C-D).

**Figure 1:**
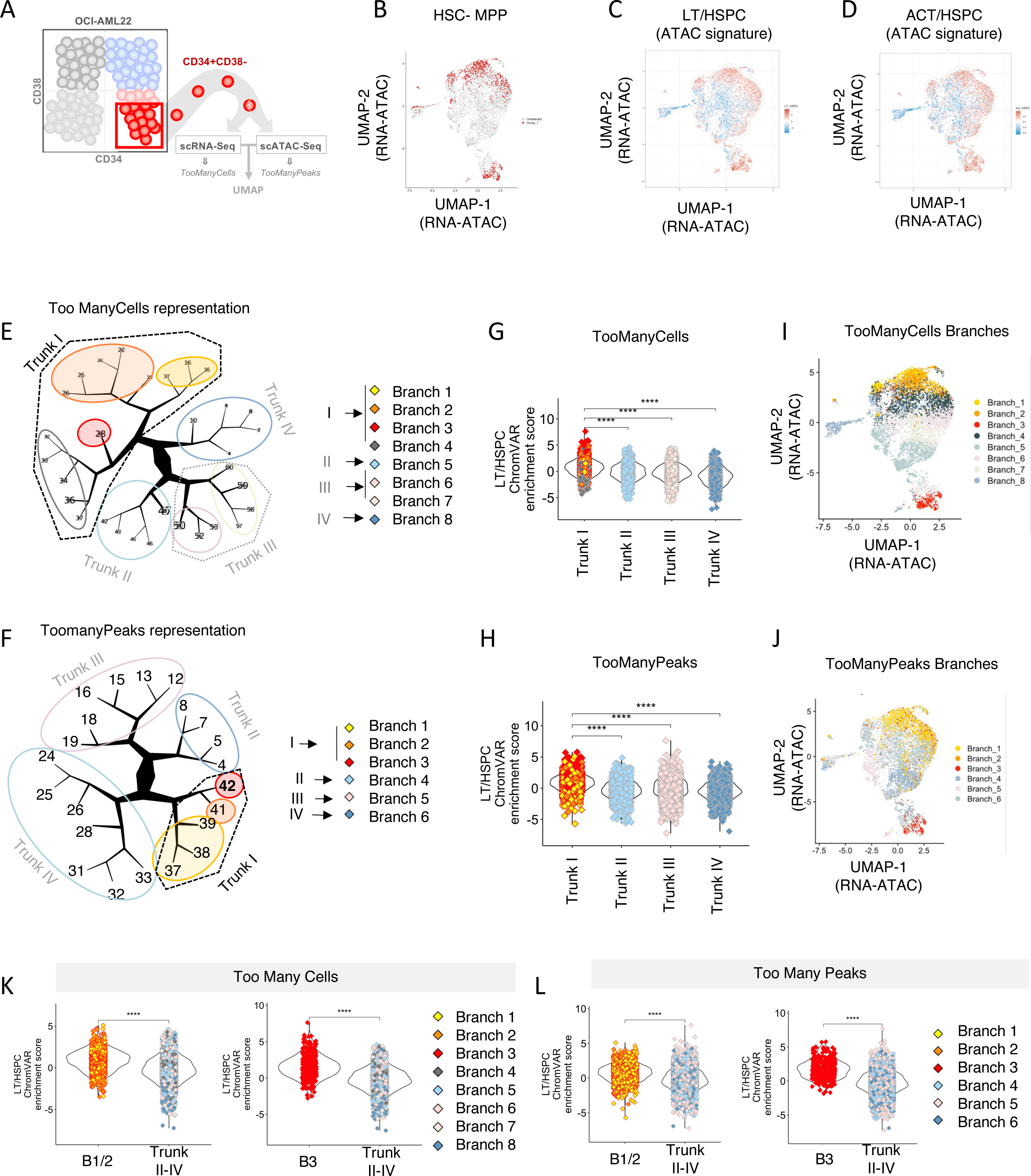
Single cell multiome analysis captures heterogeneity within the OCI-AML22 CD34+CD38-cell fraction. **A.** Experimental design. **B-D.** UMAP representation based on both RNA-Seq and ATAC-Seq for each of the CD34+CD38-OCI-AML22 cells that passed QC. Cells that mapped into the HSC-MPP group from Supplementary Figure S1 are colored in red (B) Cells are colored based on the z-score value for the LT/HSPC epigenetic signature^42^ (C) or the z-score value for the ACT/HSPC epigenetic signature from^42^ (D). **E.** TooManyCells representation^43^ using RNA-Seq data from CD34+CD38-OCI-AML22 single cells. **F.** TooManyPeak representation^44^ using ATAC-Seq data from CD34+CD38-OCI-AML22 single cells. **G.** Enrichment scores for the LT/HSPC signature^42^ are plotted for OCI-AML22 CD34+CD38-cells across the TooManyCells branches as indicated. Individual cells are colored based on their branch identity in the TooManyCells Tree (see Figure. 1E). **H.** Enrichment scores for the LT/HSPC signature^42^ are plotted for OCI-AML22 CD34+CD38-cells across the TooManyPeaks branches as indicated. Cells are colored based on their branch identity in the ToomanyPeaks Tree (see Figure 1F). **I-J.** UMAP representation based on both RNA-Seq and ATAC-Seq for each of the CD34+CD38-OCI-AML22 cells that passed QC. Cells are colored based on their branch identity in the TooManycells representation **(I**) or based on their branch identity in the ToomanyPeaks representation (**J**). **K.** Enrichment scores for the LT/HSPC signature^42^ are plotted for OCI-AML22 CD34+CD38-cells across the TooManyCells branches as indicated. Cells are colored based on their Branch identity in the TooManycells representation shown Figure 1F. **L.** Enrichment scores for the LT/HSPC signature^42^ are plotted for OCI-AML22 CD34+CD38-cells across the TooManyPeaks branches as indicated. Cells are colored based on the branch identity in the TooManyPeaks representation.

To understand the stem cell heterogeneity within the OCI-AML22 LSC fraction, we applied the TooManyCells^43^ and TooManyPeaks^44^ algorithms, which build branching trees comprising cells sharing transcriptomic or epigenetic features, respectively. Each algorithm produced a diagram with 4 trunks (Figure 1E-F), independently suggesting the existence of 4 distinct cell populations. TrunkI on both diagrams contained cells most enriched for the LT/HSPC (Figure 1G-H) or the ACT/HSPC epigenetic signatures (Supplementary Figure 4A and Supplementary Figure 5A) and most depleted for epigenetic signatures of committed populations such as monocytes or granulocytes (Supplementary Figure 4A and Supplementary Figure 5A). Interestingly, this stem-like TrunkI could be further split into multiple branches, using TooManyCells (Figure 1E) or TooManyPeaks (Figure 1F), suggesting the existence of epigenetically and transcriptionally distinct LSC subgroups making up TrunkI. Labelling each cell in the UMAP representation from Figure 1B-D based on their designated branch showed a clear allocation of individual branches associated with stemness (Branch1-3) to two distinct regions regardless of the algorithm (TooManyCells or TooManyPeaks) used (Figure 1I-J). Cells represented in Branch 1 and Branch 2 from both trees (hereafter termed B1/2-LSC) mapped to one region, while cells belonging to Branch 3 on both trees (hereafter termed B3-LSC) mapped to another distinct region (Figure 1I and Figure 1J). Both B1/2 LSC and B3 LSC branches were more enriched for the LT/HSPC epigenetic signature compared to cells that are part of the other trunks (Trunk II-IV) as identified by either TooManyCells or TooManyPeaks (Figures 1K,L), establishing their stem cell identity. Overall, these results demonstrate that two transcriptionally and epigenetically distinct LSC populations (B1/2-LSC and B3-LSC) coexist within the OCI-AML22 CD34+CD38-fraction.

### B1/2 and B3 LSC signatures are enriched in LSC+ fractions from AML patient samples and co-exist within individual patients

To determine the clinical relevance of LSC identified in OCI-AML22, we used a cohort of 73 AML patient samples that have been sorted based on CD34 and CD38 expression, where each fraction was functionally assessed for their ability to engraft NSG mice^2^. GSVA using the list of genes positively upregulated in TrunkI compared to the other Trunks (II-IV) showed that they were enriched in functionally-validated LSC-containing fractions (LSC+, 138 fractions) compared to LSC-depleted fractions (LSC-, 84 fractions)^2^ (Supplementary Figure 6A). In addition, GSVA enrichment scores correlated with LSC frequency (Supplementary Figure 6B). We then investigated whether B1/2-LSC and B3-LSC signatures could be detected within AML patient samples. To this end, we generated lists of genes that were significantly upregulated in B1/2 versus B3-LSCs and vice-versa (B1/2 and B3 signatures, respectively). GSVA scores for B1/2 and B3 signatures calculated across the 138 LSC-containing (LSC+) fractions^2^, enabled clustering these fractions into 4 clusters. One cluster was not enriched for either B1/2 or B3 signatures (Non-enriched LSC+ cluster); two clusters were enriched exclusively for either the B1/2 signature or the B3 signature (single clusters: B1/2 LSC+ cluster or B3 LSC+ cluster), and another cluster was enriched for both (Mixed LSC+ cluster) (Figure 2A). Most patient samples contained at least one LSC+ fraction enriched for B1/2 or B3 signatures (n=60/73; 81%), underscoring the broad clinical relevance of those signatures. To determine whether fractions were clustered based on intra-patient differences or if LSC+ fractions belonging to the same patient could be found across different clusters, and thus uncover heterogeneity within the LSC compartment of individual patients, we focused on patient samples with more than one LSC+ fraction (n=42/73 patients). We were able to uncover 3 groups of patient samples, based on the combinations of LSC+ fractions in each individual sample. A minority of patient samples presented only LSC+ fractions from the non-enriched cluster (n=4/42; 9.5% - Supplementary Figure 7A). The majority of patient samples presented LSC+ fractions belonging to one of the single clusters (either B1/2 or B3 clusters) (n=29/42; 69%- Figure 2B-C). Interestingly, a third group of patient samples (n=9/42; 21%) exhibited a more complex composition of LSC+, with a combination of B1/2 and B3 signatures either present in separate fractions (Figure 2D) or in the same fraction (Figure 2E-H). Taken together, these data demonstrate that the two LSC subtypes (B1/2 and B3) we identified in OCI-AML22 are found in the majority of primary AML samples and can co-exist, in varying combinations, within individual patients.

**Figure 2:**
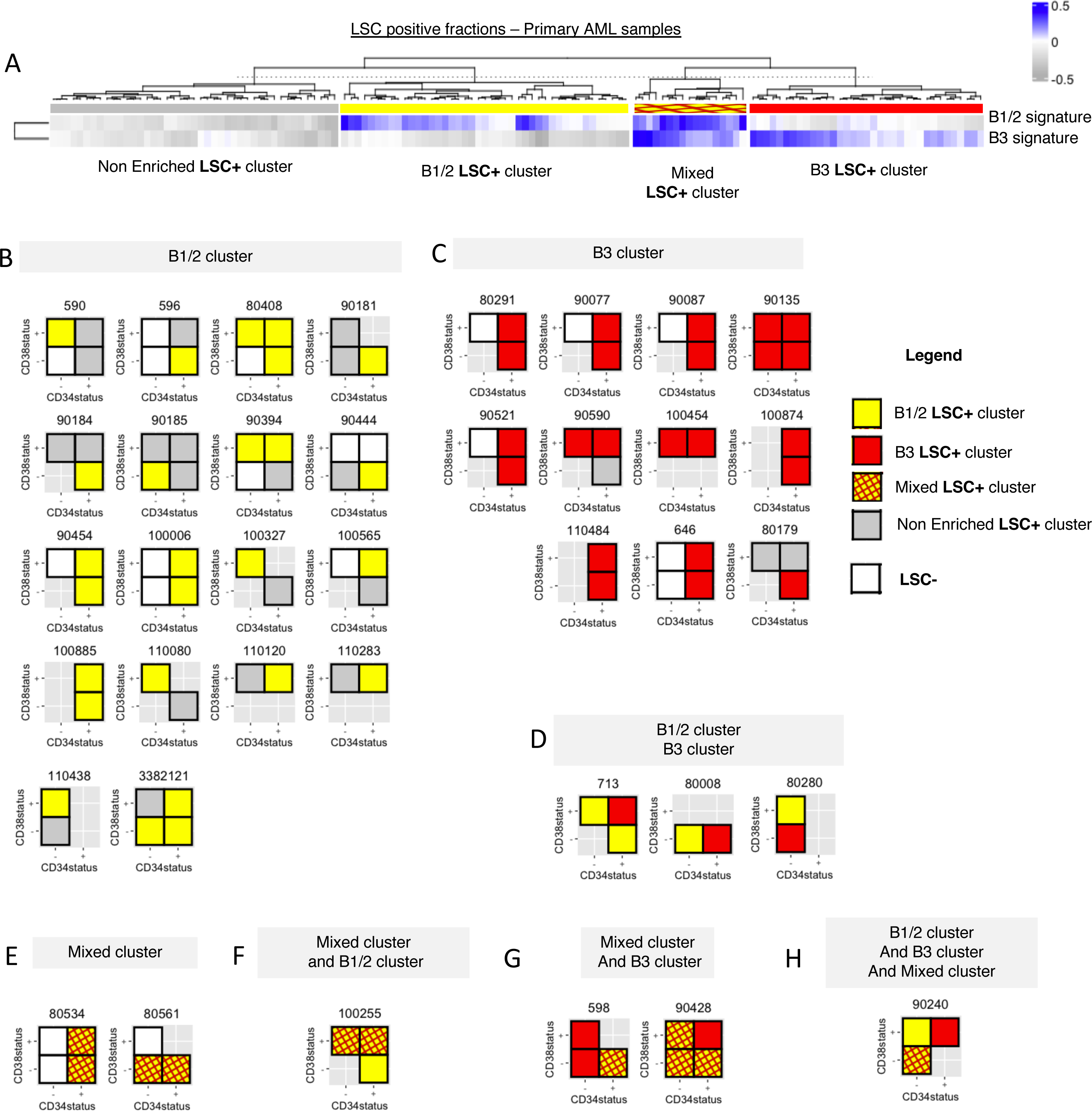
B1/2 and B3 LSC signatures can be identified in LSC+ fractions from primary AML patient samples and co-exist within an individual LSC+ fraction. **A.** GSVA was performed for B1/2 or B3 signatures across functionally assessed LSC-positive fractions from primary AML samples^2^. Scores for each signatures are plotted and were used to perform hierarchical clustering using Complexe heatmap. **B-H.** Functionally assessed LSC positive fractions for each indicated patient sample are represented in boxes colored based on the cluster each fraction was allocated in Figure 2A. LSC negative fractions are indicated in white. Not assessed fractions are not represented. Patients with fractions belonging to the B1/2 cluster from Figure 2A (B), Patients with fractions belonging to the B3 cluster from FIgure 2A (C), Patients with fractions belonging to cluster B1/2 or cluster B3 from figure 2A (D), Patients with fractions belonging to the Mixed cluster (E), Patients with fractions belonging to the Mixed cluster and the B1/2 cluster (F), Patients with fractions belonging to the Mixed cluster and the B3 cluster (G), Patient with fractions belonging to the 3 clusters enriched with either B1/2 or B3 signatures (H).

### Single cell assay captures distinct colony forming and differentiation potentials within the CD34+CD38-OCI-AML22 fraction

Having identified two groups of LSC based on transcriptional and epigenetic features, we next investigated whether there was functional heterogeneity in the LSC compartment. To this end, we first developed an *in vitro* assay to analyze the progeny produced by single OCI-AML22 CD34+CD38-cells deposited into 96 well plates pre-seeded with MS5 stromal cells (Figure 3A)^45^. Approximately half of the seeded CD34+CD38-cells were able to generate a colony (Figure 3B). We distinguished two types of clonogenic outputs (Figure 3C-H). The first type was characterized by smaller colonies retaining CD34+ expression on >80% of cells. In contrast, the second type was characterized by larger colonies with <80% CD34+ cells. Both colony types exhibited similar expansion of the CD34+CD38-population from the original single seeded cell despite their differential capacity to generate large numbers of downstream progenitors (Figure 3E-H). Collectively, these data point to the existence of two clonogenic cell types within the OCI-AML22 CD34+CD38-fraction, each able to expand this fraction but differing in their differentiation potential.

**Figure 3:**
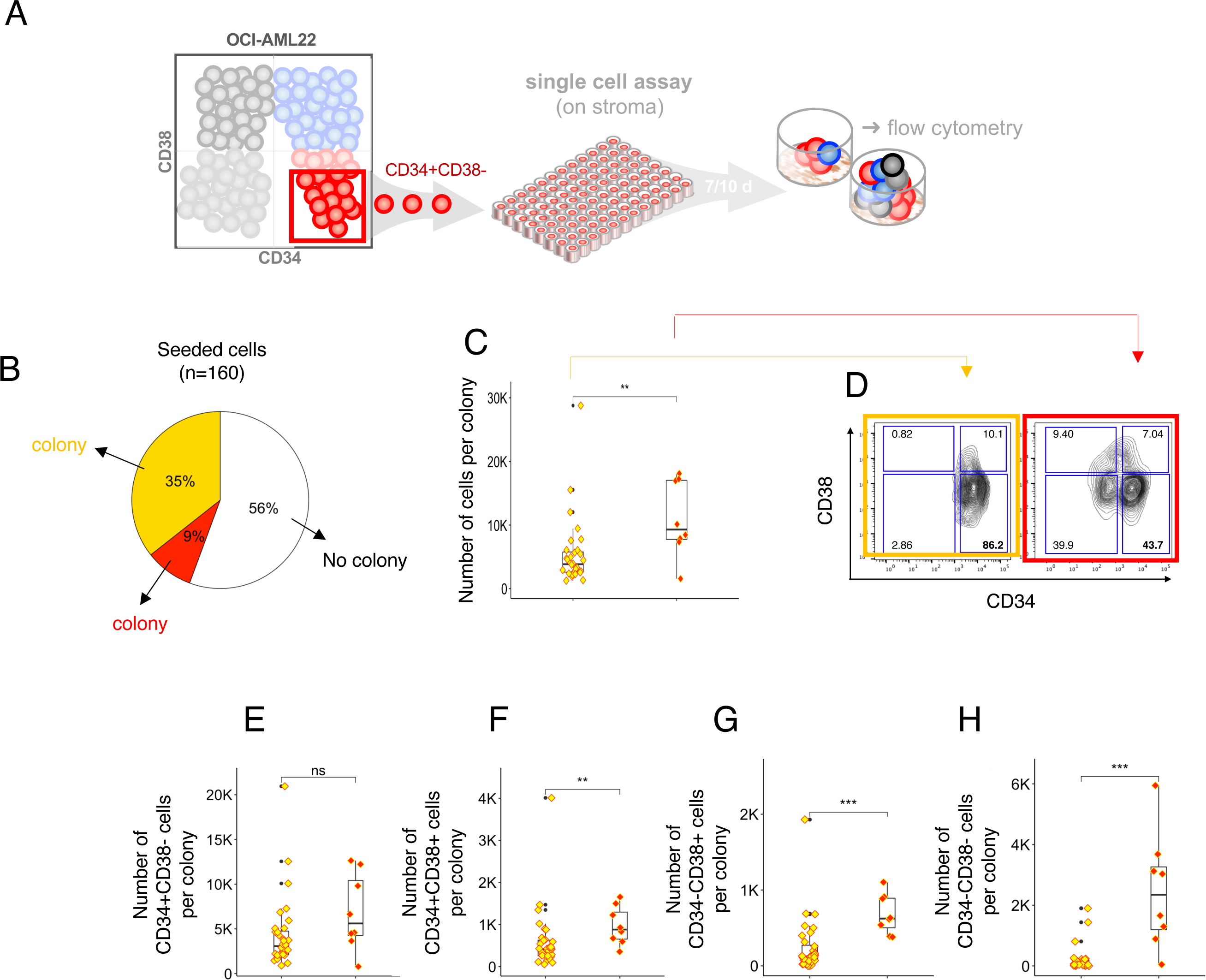
Single cells assay captures distinct colony formation and differentiation potentials within the CD34+CD38- OCI-AML22 fraction. **A**. Experimental scheme. **B**. Pie chart indicating the percentage of CD34+CD38- OCIAML22 cells that do not generate colonies (depicted in white) or generate a colonie (depicted in yellow or red depending on the type of colonies). **C**. For each colony type depicted in yellow or red in Figure 3B, the absolute number of total cells is plotted in the matching color (yellow: left; red: right). **D**. Extreme immunophenotypic profiles of the 2 types of colonies identified from single cell in vitro assay and depicted in yellow or red in Figure 3B. **E-H**. For each colony type depicted in yellow or red in Figure 3B, the absolute number of CD34+CD38- cells (E), CD34+CD38+ cells (F), CD34- CD38+ cells (G) or CD34-CD38-cells (H) generated from each single CD34+CD38-OCI-AML22 cells co-cultured with MS5 is depicted in the matching colors (yellow: left; red: right).

### Xenotransplantation assays reveal the existence of LSC with distinct repopulation kinetics and differentiation potentials

To determine if the functional heterogeneity we found *in vitro* could also be identified and functionally validated *in vivo*, we xenotransplanted OCI-AML22 CD34+CD38- cells at limiting dilution to assess the repopulation potential of individual LSCs after 8 and 12 weeks (Figure 4A). Interestingly, more grafts were detectable both at the limiting dose (469 cells per mouse) and at the intermediate dose (1875 cells per mouse) after 12 weeks compared to 8 weeks (Figure 4B). Given that stem cell detection requires generation of a large enough graft to be measurable, our results suggest variability in the kinetics of engraftment of individual LSCs, with one LSC subtype repopulating mice faster to become detectable at 8 weeks, compared to the other. To gain a deeper understanding of the functional heterogeneity observed *in vivo* within the pool of OCI-AML22 LSC, we performed dimensionality reduction using UMAP of experimental parameters including the number of cells injected, engraftment level as a surrogate for repopulation capacity, and the proportion of cells expressing CD34 and CD38 as a surrogate for differentiation capacity. The UMAP analysis revealed the existence of 2 clusters (Figure 4C). Interestingly, despite the fact that the number of injected cells per mouse was similar (Figure 4D), the 2 clusters differed in engraftment level (Figure 4E) and the CD34+ expression profile of the resulting graft (Figure 4F-G). Collectively, these data suggest the existence of 2 subtypes of functional LSC in OCI-AML22: one subtype that repopulates slowly to generates grafts that are only detectable after 12 weeks, while a second subtype that repopulates mice more rapidly and thereby, producing sufficient progeny for the graft to be detected as early as 8 weeks. These distinct differentiation and expansion capacities are reminiscent of the functional differences we observed *in vitro*.

**Figure 4:**
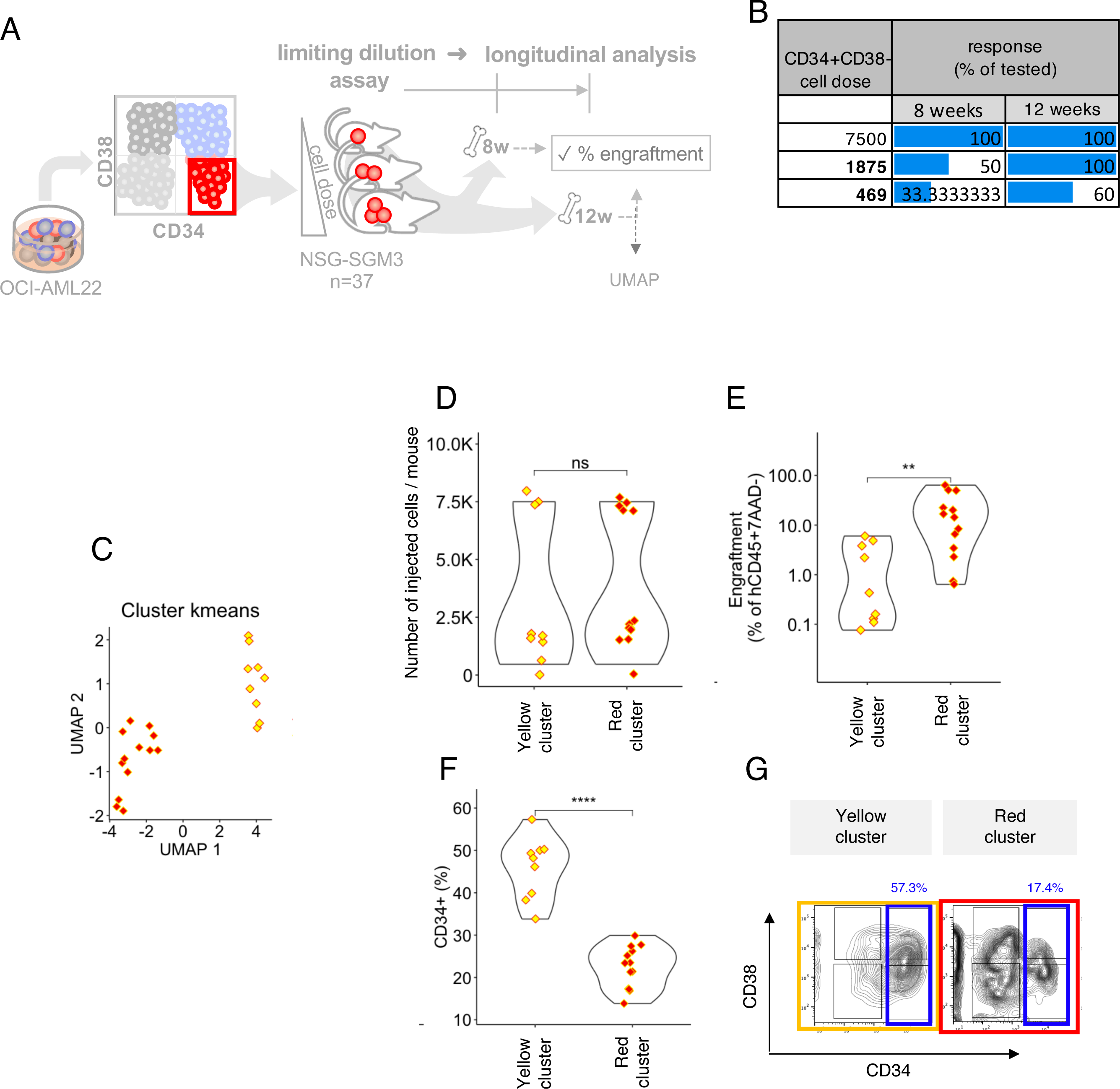
In vivo functional deconvolution reveals the existence of LSC with distinct repopulation kinetics and differentiation potential. **A.** Schematic representation of the experiment. NSG-SGM3 mice were injected intrafemorally at limiting dilution with multiple cell doses of sorted CD34+CD38- OCI-AML22 cells. Engraftment was assessed 8 and 12 weeks after injection. **B**. The percentage of mice for which engraftment could be detected at 8 weeks or 12 weeks, after injection of the indicated OCI-AML22 CD34+CD38- cell dose sorted from a bulk culture expanded for 3-4 months in vitro is plotted. **C.** UMAP-based clustering of engraftment parameters derived from non-injected bone marrow (BM) and injected femur (RF) (parameters: total human cell counts, %CD34 expression, %engraftment level, number of injected cells per mice) of NSG-M3S mice 12 weeks after injection CD34+CD38- OCIAML22 cells per mouse using doses from Figure 4B. **D**. Statistical analysis for each cluster from Figure 4C, showing the number of injected cells (D), the engraftment level (E), the percentage of CD34+ cells in the generated grafts (F). **H**. Representative immunophenotypic profiles of grafts based on CD34 and CD38 cell surface expression (hCD45+ subgated cells are shown) for the clusters identified in Figure 4C.

### CD112 enables prospective isolation of distinct LSC subsets linked to different cell cycle states

We previously characterized two populations of normal LT-HSC that can be segregated based on expression of CD112 (CD112High and CD112Low) and that exhibit distinct repopulation kinetics linked to differences in cell cycle state^11^. In this prior study, we generated a chromatin accessibility signature consisting of peaks more accessible in the CD112High population compared to the CD112Low population (CD112High signature). To assess whether B1/2 and B3 LSC from OCI-AML22 could be distinguished by CD112 expression, we performed ChromVAR enrichment for the CD112High signature and found that B3-LSC were more enriched for this signature compared to B1/2-LSC (Figure 5A). To examine whether the repopulation kinetics of the two LSC subtypes we identified could be explained by distinct cycling capacities, we calculated cell cycle phase scores using canonical markers of G2M or S phases^46^. Cells part of TrunkI from TooManyCells (Figure 1) were the most enriched in G0/G1, suggesting they are not actively cycling, compared to cells that were found in the other Trunks (Figure 5B). This is consistent with the fact that LSC are less proliferative than their downstream progeny^47^. To further discriminate cycling abilities within the less proliferative TrunkI cell population, we performed GSEA using 3 independent lists of genes known to be more expressed in cycling cells (G2M) compared to quiescent cells (G0/G1): LSPC-Cycle top 250 genes^33^, Tirosh Cycle^46^ and the Rehman Colon Diapause Down^48^ signatures. B3-CD112High LSC were more enriched for these cycling signatures compared to B1/2-CD112Low LSC (Figure 5C), suggesting that the former corresponds to the more proliferative, rapidly repopulating cells identified in our *in vitro* and *in vivo* functional assays. To determine whether B1/2 and B3 LSCs can be separated based on CD112 cell surface expression, CD34+CD38- OCI-AML22 cells were sorted based on CD112 expression (CD112High, top 20%; CD112Low, bottom 20%; Figure 5D). Cell cycle was assessed using classic markers (Ki67 and Hoechst) known to distinguish between G0 (Hoechst^low^ Ki67-) and G1 cells (Hoechst^low^Ki67+). Additionally we also measured CDK6 expression, a marker of G0 exit that can further stratify G0 cells into deeply quiescent cells (CDK6-Ki67-) versus cells primed for more rapid G0 exit (CDK6+KI67-)^7,11^. The CD112Low fraction contained a higher proportion of Hoechst^low^ Ki67- and Ki67-CDK6-cells compared to the CD112High fraction (Figure 5E-F). To extend our findings to primary samples, we examined functionally defined LSC+ fractions from two AML patient samples. T, the CD112Low population in both cases contained a higher proportion of cells in G0 (Hoechst^low^ Ki67-) and a greater proportion of CDK6-Ki67-deeply quiescent cells (Figure 5G,H) compared to CD112High cells. Together, these data demonstrate that CD112 expression provides a tool for prospective enrichment of two LSC subtypes displaying distinct cell cycle states.

**Figure 5:**
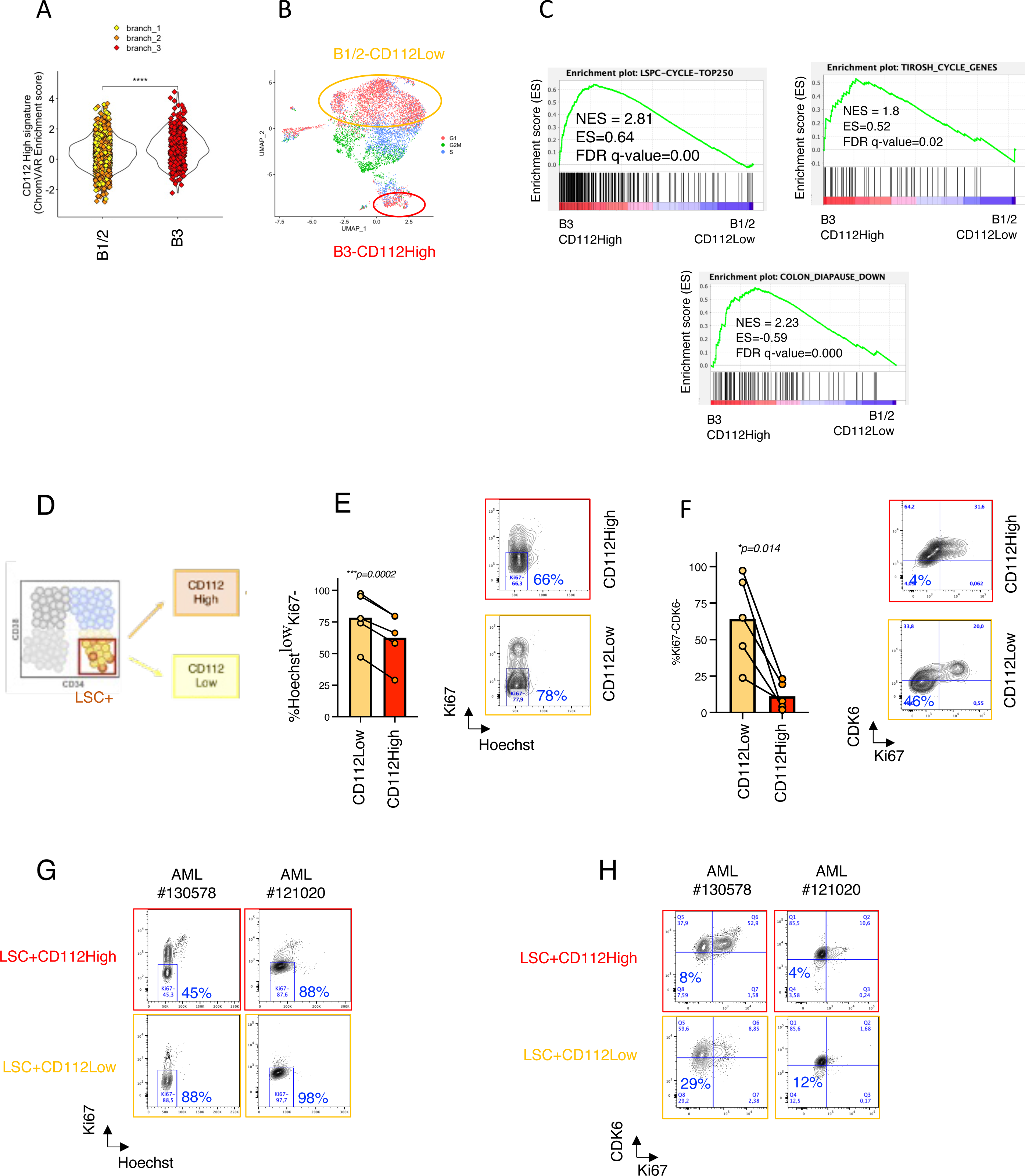
CD112 enables prospective isolation of distinct LSC subsets with different cell cycle state. **A.** ChromVAR enrichment score for the CD112-High epigenetic signature^11^ in the indicated populations from TooManyCells. **B.** UMAP representation based on both RNA-Seq and ATAC-Seq for each of the CD34+CD38- OCI-AML22 cells that passed QC. Cells are colored based on the cell cycle phase they were allocated to using Seurat cell cycle signature^46^. **C.** GSEA across B3-CD112High and B1/2-CD112Low for LSPC-Cycle Top 250 signature^33^, Tirosh cell cycle signature^46^ and the Diapause Down in colon cancer signature^48^. **D.** Experimental scheme. **E-F.** OCI-AML22 CD34+CD38- fraction was sorted based on CD112 expression level then stained for CDK6, Ki-67 and Hoechst to determine the percentage of cells in G0 (Hoechst^Low^Ki67-) (E) or the deepest quiescent cells Ki67-CDK6-cells (F). Representative FACS plots are disclosed for each condition on the right (E-F). **G-H.** The LSC+fraction from functionally assessed primary AML samples was sorted based on CD112 expression level (top 20% :CD112-High, bottom 20% :CD112-Low) then stained for CDK6, Ki67 and Hoechst to determine the percentage of cells in G0 (Hoechst^Low^Ki67-)(G) or the deepest quiescent cells in G0 (Ki67-CDK6-) cells (H). FACS plots are disclosed for each condition for each patient sample (G-H).

### Prospectively isolated LSC subtypes preserve their distinct differentiation potentials through serial repopulation

To demonstrate the link between CD112 expression, cell cycle state and repopulation ability of the two identified LSC subtypes, we injected CD112High and CD112Low fractions sorted from OCI-AML22 cells into NSG mice (Figure 6A). Although both fractions were able to engraft mice, CD112Low cells gave rise to smaller grafts compared to CD112High cells (Figure 6B) and presented a phenotype reminiscent of the two distinct LSC outputs we found from our *in vitro* and *in vivo* studies, although only 1 mouse transplanted with CD112Low cells showed engraftment over 20%. To assess long-term repopulation and differentiation capacities of the two prospectively isolated LSC populations, we isolated CD34+CD38- cells (regardless of CD112 expression) from primary grafts generated by either CD112Low or CD112High cells and injected them into secondary NSG-SGM3 mice (Figure 6D). Secondary grafts generated by cells harvested from CD112Low primary grafts contained a higher proportion of CD34+ cells (Figure 6D-E) compared to the secondary grafts generated by cells harvested from CD112High primary grafts. Collectively, these findings establish the existence of at least two LSC subtypes that can be prospectively isolated based on CD112 expression and exhibit distinct repopulation kinetics and differentiation potentials that are retained through serial transplantation (Figure 6F).

**Figure 6:**
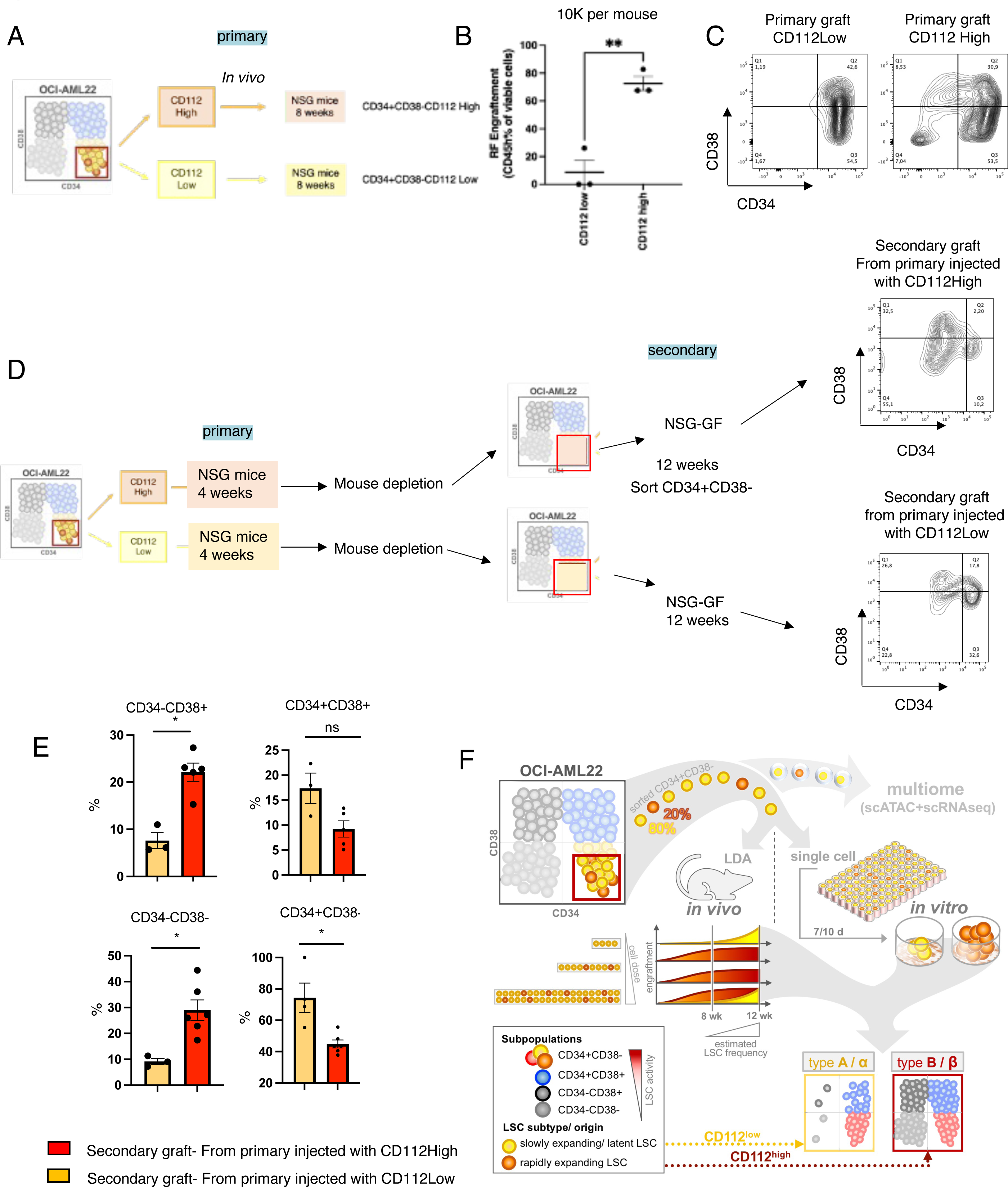
Prospectively isolated LSC subtypes preserve their distinct differentiation potentials throughout serial repopulation assays. **A.** Experimental in vivo scheme. **B.** Engraftment level 8 weeks after injection of 10 000 OCI-AML22 cells in NSG mice sorted as indicated on Figure 6A. **C.** Experimental in vivo strategy for secondary experiment and representative FACS plots for each condition based on CD34 and CD38 among the CD45+7AAD- engrafted population. **E.** Percentage of CD34/CD38 cells of secondary grafts generated after injection of 10 000 CD34+CD38-cells CD112 High or CD112 Low, as indicated in Figure 7D. Mann Whitney test, n=9 mice. **F.** Summary scheme.

## Discussion

Integrating a panel of functional *in vitro* and xenograft assays, with single cell multiomics, we report the existence of two LSC subtypes with distinct epigenetic and transcriptional signatures. These subtypes are associated with distinct functional properties including differences in their level of quiescence and repopulation capacity that are retained through serial transplantation. Importantly, these two LSC subtypes could be prospectively purified based on CD112 cell surface expression, demonstrating that the two subtypes are distinct stem cell entities and not a result of stochastic cell cycle fluctuations within a homogenous LSC population. Our findings complement studies that have shown the existence of genetic subclones across primary AML samples^17,49^, as we have also recently demonstrated for the OCI-AML22 model^34^, thereby bringing additional insight into LSC properties at the transcriptomic and epigenetic level.

The identification of two novel functional LSC subsets in AML that differ at the transcriptomic and epigenetic level is of particular interest as it is widely considered that transcriptional and epigenetic features are more amenable to pharmacological perturbation^50^ than mutationally-driven alterations^15,51–59^. Given their specific signatures, the identified LSC subtypes may exhibit different responses to individual therapies. We previously reported a cellular hierarchy classification for AML that represents a powerful approach to predict drug response^33^. However, in this prior study we did not link any specific LSC subtype to a distinct hierarchy classification nor investigated the possibility for patients to present multiple LSC types, as no markers had been identified that could reliably separate functional LSC subsets. The ability to exploit CD112 as a means for prospective enrichment of distinct LSC subsets now provides a crucial tool for refining our understanding of the LSC compartment, including identifying specific vulnerabilities of all the LSC subsets that might exist in an individual patient.

The OCI-AML22 model was central to our study as we were able to undertake deep characterization and purification of the distinct LSC fractions. The LSC signatures derived from the distinct LSC subtypes found in OCI-AML22 were also captured within LSC fractions extracted across a large cohort of AML patient samples and congruent with our model can co-exist within the same AML patient sample. Collectively, these findings highlight the utility of the OCI-AML22 model to unlock LSC properties at a functional and mechanistic level and point to the broad biological relevance of our findings. Thus, we envision that the OCI-AML22 model, combined with the potential to derive genetic sublines or increase clonal diversity via CRISPR editing and lentiviral technologies^34^, will serve as a platform for interrogating inter- and intra-patient LSC heterogeneity and extracting shared intrinsic and even clone-dependent LSC vulnerabilities. Altogether, our study sets the stage to unlock new insights into the mechanisms governing stemness in leukemia and beyond.

## Acknowledgements

J.E.D is supported by funds from the: Princess Margaret Cancer Centre Foundation, Ontario Institute for Cancer Research through funding provided by the Government of Ontario, Canadian Institutes for Health Research (RN380110 - 409786), Canadian Cancer Society (grant #703212 (end date 2019), #706662 (end date 2025), Terry Fox New Frontiers Program Project Grant (Project# 1106), a Canada Research Chair, Princess Margaret Cancer Centre, The Princess Margaret Cancer Foundation, and Ontario Ministry of Health. HB was supported by the Helena Lam Fellowship in Cancer Research Fund., the Leukemia Tissue Bank at Princess Margaret Cancer Centre/ University Health Network as the source of primary sample blood. We thank Suraj Bansal, Sayyam Shah and Andy Zeng for helpful discussions and members of John Dick lab for reading the manuscript.

## Author contributions

H.B. conceived the study, performed research, performed bioinformatic analysis and wrote the manuscript. A.M. performed bioinformatic analysis and edited the manuscript, C.A. Performed research. M.L. provided funding. J.W. edited the manuscript and provided conceptual input. K.B.K. performed research, provided conceptual input and wrote the manuscript. J.E.D. edited the manuscript, provided conceptual input, secured fundings and supervised the study.

## Material and Methods

### OCI-AML22 culturing

cells were expanded in culture in the following medium, with final concentrations as indicated : X-VIVO 10 (Lonza, BE04-380Q) supplemented with 20% BIT 9500 Serum Substitute (StemCell technologies, 09500), 1x Glutamax Supplement (Thermo Fisher Scientific, 35050061), Primocin 0.1 mg/ml (invivogen), SCF (200 ng/ml;Miltenyi Biotech, 130-096-696), IL3 (20 ng/ml;Miltenyi Biotech, 130-095-069), TPO (20 ng/ml; Peprotech, 300-18), FLT3L (40 ng/ml; Peprotech, 300-19 B), IL6 (10 ng/ml; Miltenyi Biotech, 130-093-934), G-CSF (10 ng/ml; Miltenyi Biotech, 130-093-861). Cells were maintained at a density of 0.8×10^6^ cells/ml and passaged every 3-5 days, in a 96 well flat bottom plate. The CD34+CD38- fraction or CD34+ fraction was regularly sorted to serially expand the cells.

### Xenotranplantation

All animals used for this study were treated in accordance with institutional guidelines. Female and male NSG mice (NOD.Cg PrkdcscidIl2rgtm1Wjl/SzJ; The Jackson Laboratory) or male NSG-SGM3 mice as indicated in figure legends were irradiated with 250 rads the day before intrafemoral injection. Bulk cells expanded in vitro for an average of 4 months were used throughout this study unless specified otherwise and are considered the OCI-AML22 model. Fractions were obtained from expanded cells in vitro after an average of 4 months, then sorted according to their CD34 and CD38 expression, using AnnexinV-FITC (BD) and 7-AAD to exclude apoptotic and dead cells. The number of injected cells per experiment is indicated in the figure legends. For lentiviral transduction experiment, cells were transduced with the ATF4 reporter as described in ^60^ in their standard culture medium overnight at a density of 0.8×10^6^ cells/ml, washed and injected the day after. Mice were euthanized and the injected femur (right femur /RF) and the non-injected left femur as a surrogate for engraftment in the bone marrow (BM) were flushed separately in MEMalpha with 10% FBS. Engraftment level was assessed by flow cytometry on a FACSCelesta instrument (BD) with the following antibodies: mCD45-FITC or hCD45-FITC, hCD45-APC-Cy7, hCD34-PE, hCD38-PE/Dazzle, AnnexinV-FITC and 7-AAD; all from BD. For secondary engraftment, human cells collected from NSG engrafted mice were sorted using 7AAD, AnnexinV-FITC, hCD45-APC-Cy7, then injected into NSG-SGM3 mice at the indicated cell dose per mice and sacrificed at 8 weeks after. The engraftment level was determined as for the primary engraftment assay.

### Multiome analysis

Raw sequencing reads were aligned to GRCh38 using cellranger-arc with default settings. Amulet and DoubletFinder were used to predict doublets from each of the scATAC and scRNA-Seq components, and predicted doublets were filtered from the dataset. Cells were further filtered based on QC metrics calculated via Signac; number of ATAC-Seq reads per cell (between 1000-100000) RNA-Seq reads per cell (1000-25000), nucleosome signal (<2) and TSS Enrichment (>1). MACS2 was used to compute a new background peak-set for all remaining cells, and UMAP performed upon the joint embedding of scRNA-Seq (following principal components analysis) and scATAC-Seq over these sites (following latent semantic indexing).

ChromVAR was used to compute the per-cell enrichment of various chromatin signatures relative to our background peak-set. TooManyPeaks (scATAC-Seq) and TooManyCells (scRNA-Seq) were performed with default settings upon the dataset to identify - due to dataset size, for TooManyPeaks the LSI reduction of scATAC was used as input in place of raw counts, omitting the first dimension, and a minimum of 100 and 200 cells per node was required when pruning results for TooManyCells and TooManyPeaks respectively. For differential expression, cells with > 5% ribosomal reads were ignored, raw counts were log-normalized, and scaled using seurat based on the top 3,000 most variable genes. scRNA-Seq counts were used in combination with the branches identified via TooManyCells as input for CIBERSORTx to create a signature matrix to deconvolute bulk samples based upon these branches. Bulk samples were deconvoluted based upon these signatures using default parameters, and normalized with the original scRNA-Seq counts.

### In vitro single cell assay

MS-5 stromal cells were kindly gifted from Dr. K. Itoh from Japan^45^. MS-5 cells were seeded into 0.2% gelatin coated 96 well-flat bottom plates (Nunc) at a density of 4-5×105 cells per well in Myelocult medium (H5100, Stem Cell Technology). On the morning, medium was changed for OCI-AML22 culturing medium (50uL/well). The CD34+CD38-OCIAML22 fraction from cultured cells (about 3-4 months) was sorted and directly plated in 96 wells plates precoated with MS5, then refilled with 50uL of fresh medium per well. 7 to 10 days after, half of each well was collected, washed and stained with hCD45-FITC, hCD34-APC-Cy7, hCD38-APC, 7-AAD, then acquired on a FACSCelesta (BD). Wells for which more than 10 cells viable cells could be detected in the gate hCD45+ was identified as a colony. To determine the percentage of CD34/CD38 within the colony, a minimum number of 300 cells in the gate hCD45+ viable cells was set in order to increase confidence in the percentage.

### CD112-based sorting and cell cycle analysis

Surface staining of OCI-AML22 cells was performed with anti-CD34-APC-Cy7, anti-CD38-PE-Cy7 and anti-CD112-APC and combined with dead cell exclusion (SytoxBlue; AnnexinV-FITC) for sorting CD34+CD38- cells into CD112High (top 20%) and CD112Low (bottom 20%) fractions on a BD Aria Fusion instrument. Cell preparation for intracellular staining of sorted cells was performed with BD Cytofix/Cytoperm according to manufacturer’s recommendations and incubated with anti-Ki67-FITC and CDK6-AF647 in PermWash solution (BD) overnight. Prior to flow cytometry analysis on a BD Fortessa instrument, cells were stained with Hoechst 33342 (1:2000; Thermo Fisher). Positive gates were set according to unstained or isotype controls and data was analyzed by FlowJo.

### Statistical analysis

GraphPad Prism or R was used. Unless specified, Mann Whitney test was performed. *<0.05, **p<0.01, ***p<0.001.

---

**Supplementary Figure S1.**
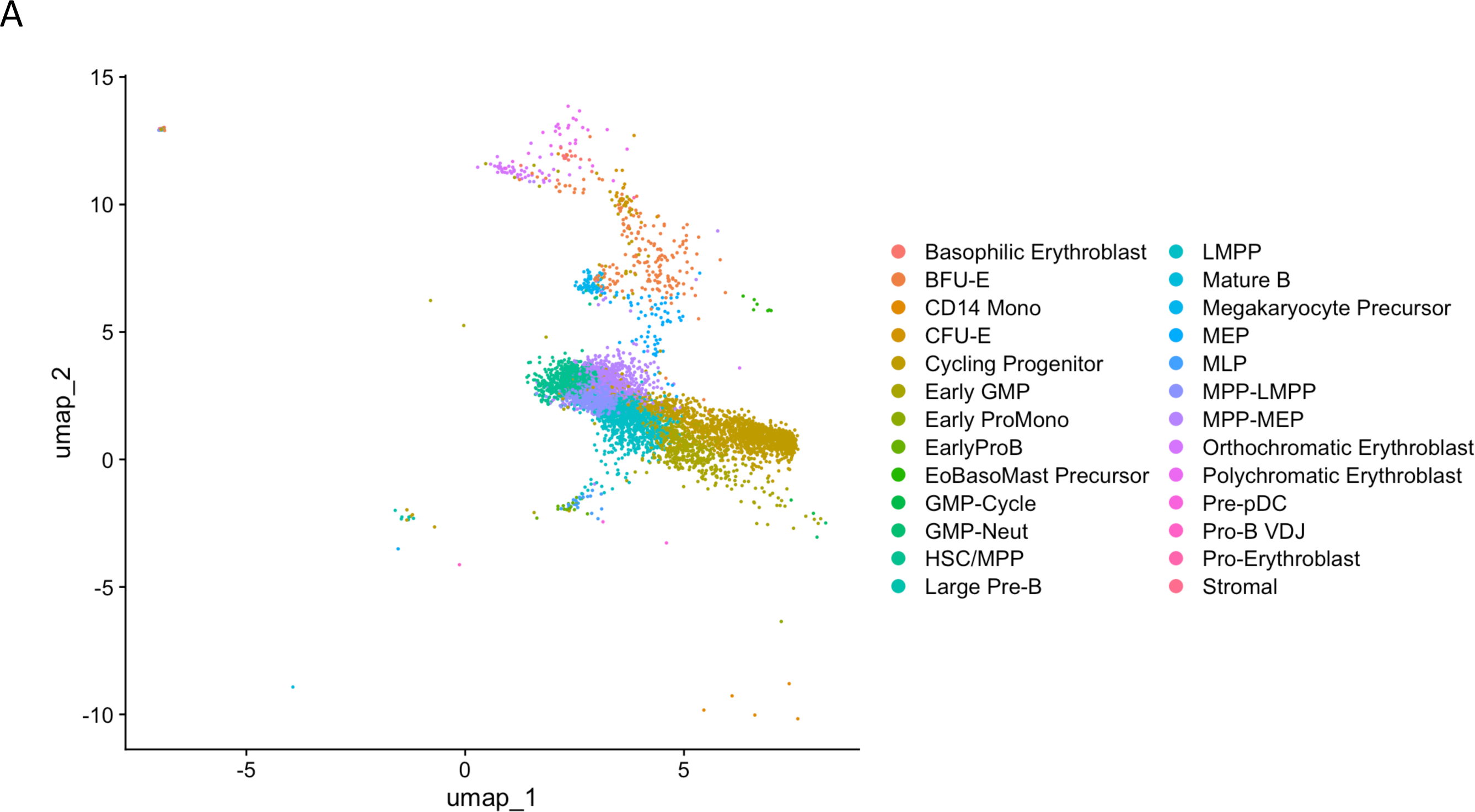
**A.** multiome paired scRNA_Seq/sc-ATAC-seq was performed on CD34+CD38- OCIAML22 cells that were then mapped onto a bone marrow reference map^41^ to label cells based on which group it colocalized with.

**Supplementary Figure S2.**
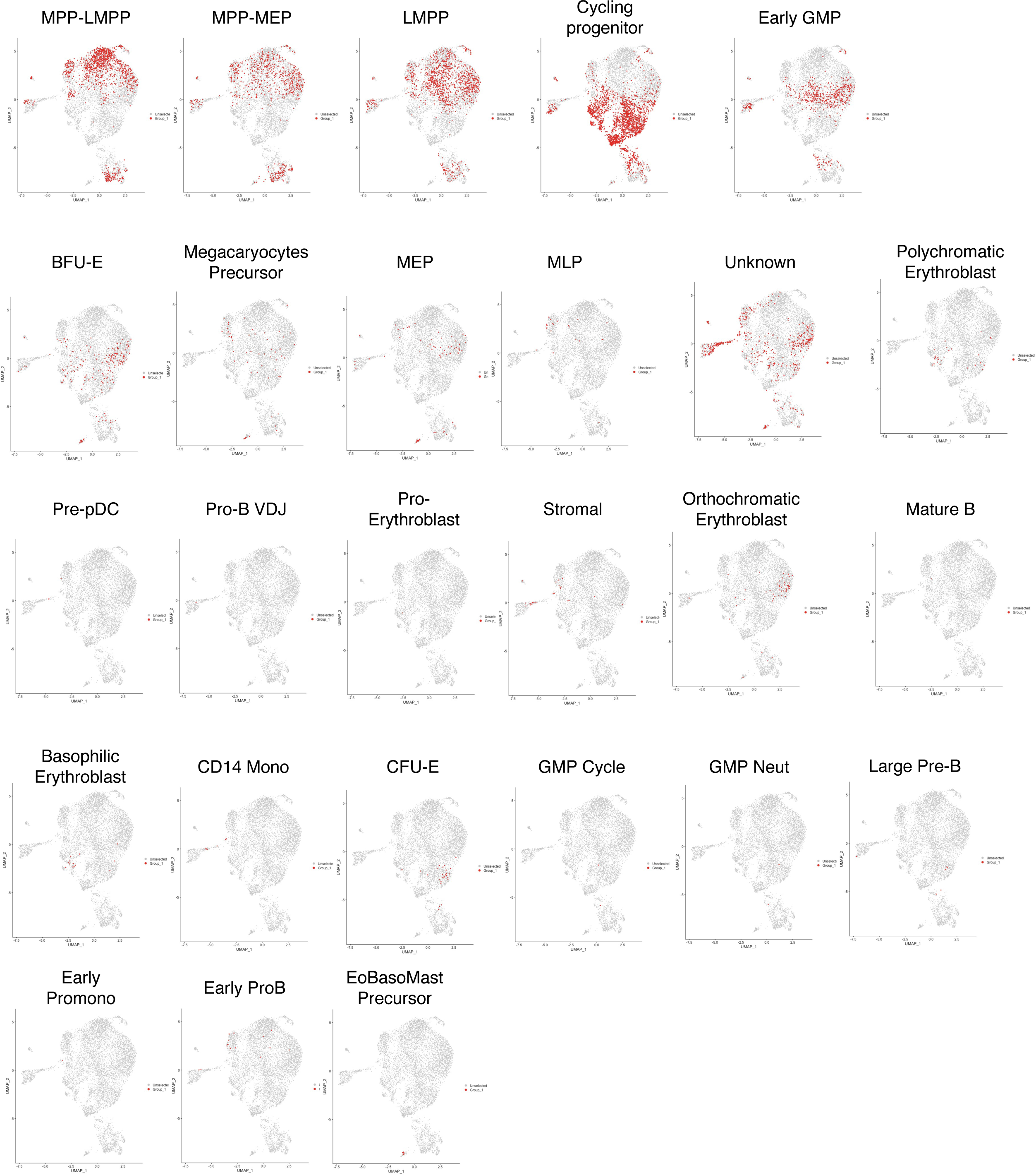
**A.** UMAP representation based on both RNA-Seq and ATAC-Seq for each of the CD34+CD38- OCI-AML22 cells that passed QC. Cells are colored based on the population they belonged to using the Reference Map^41^ (red: cells of interest, grey: others).

**Supplementary Figure S3.**
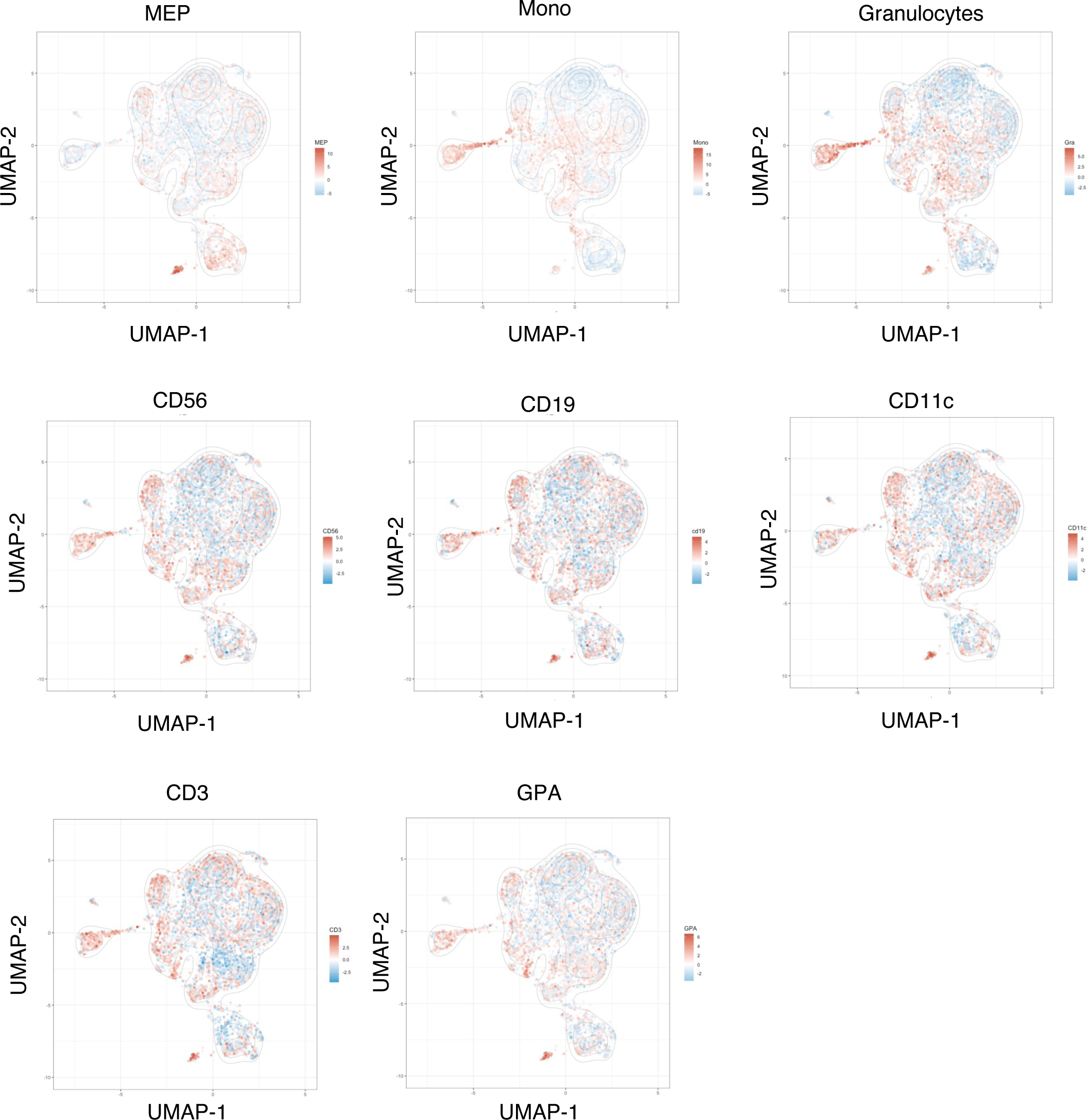
**A**. UMAP representation based on both RNA-Seq and ATAC-Seq for each of the CD34+CD38- OCI-AML22 cells that passed QC. Cells are colored based on the z-score value for the epigenetic signature from ^42^ as indicated on top of each plot.

**Supplementary Figure S4.**
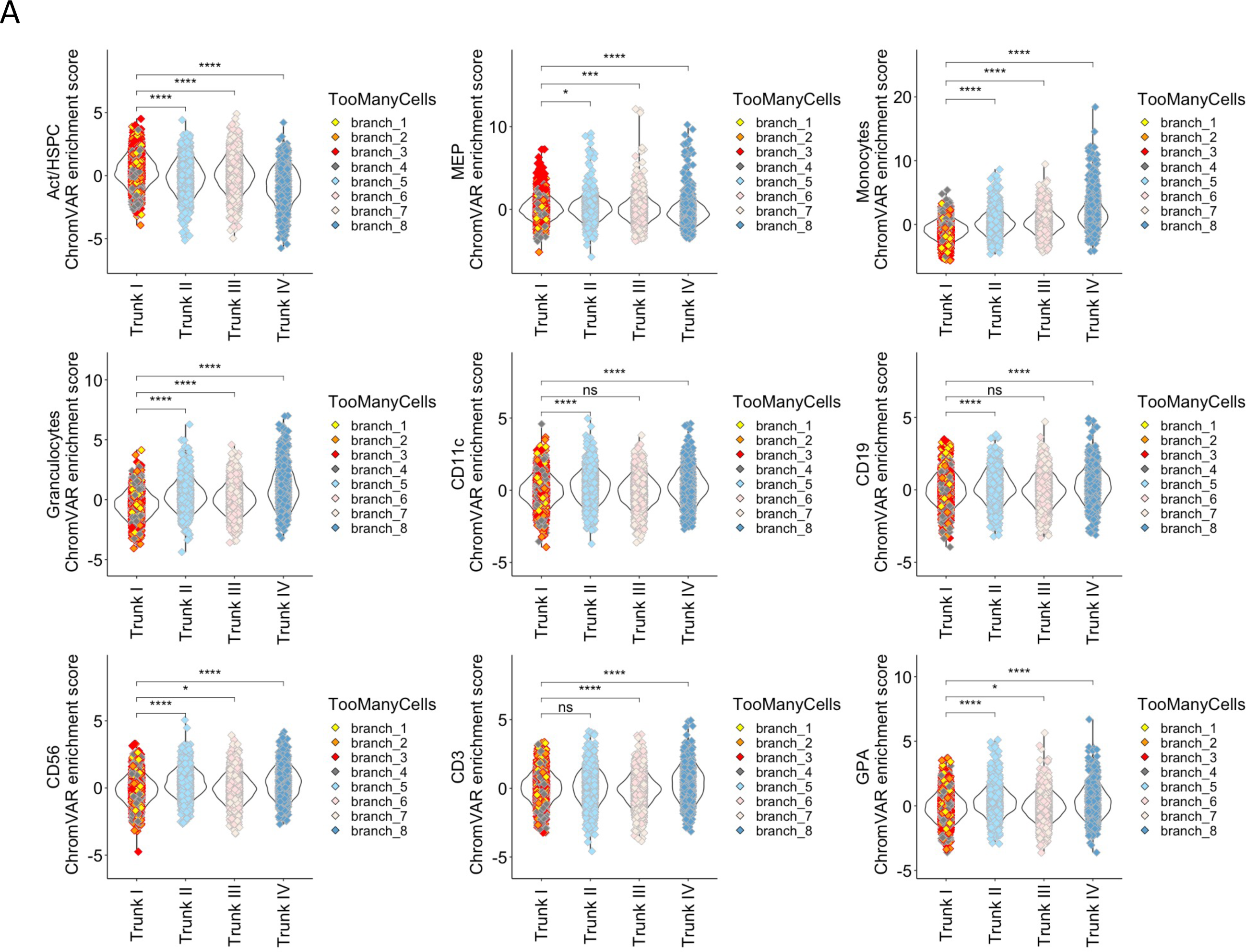
**A.** ChromVAR enrichment scores for each of the epigenetic signatures from ^42^ are disclosed for OCI-AML22 CD34+CD38- cells across the ToomanyCells branches as indicated. Cells are colored based on the Branch they were allocated to in the Toomanycells Tree from figure 1E.

**Supplementary Figure S5.**
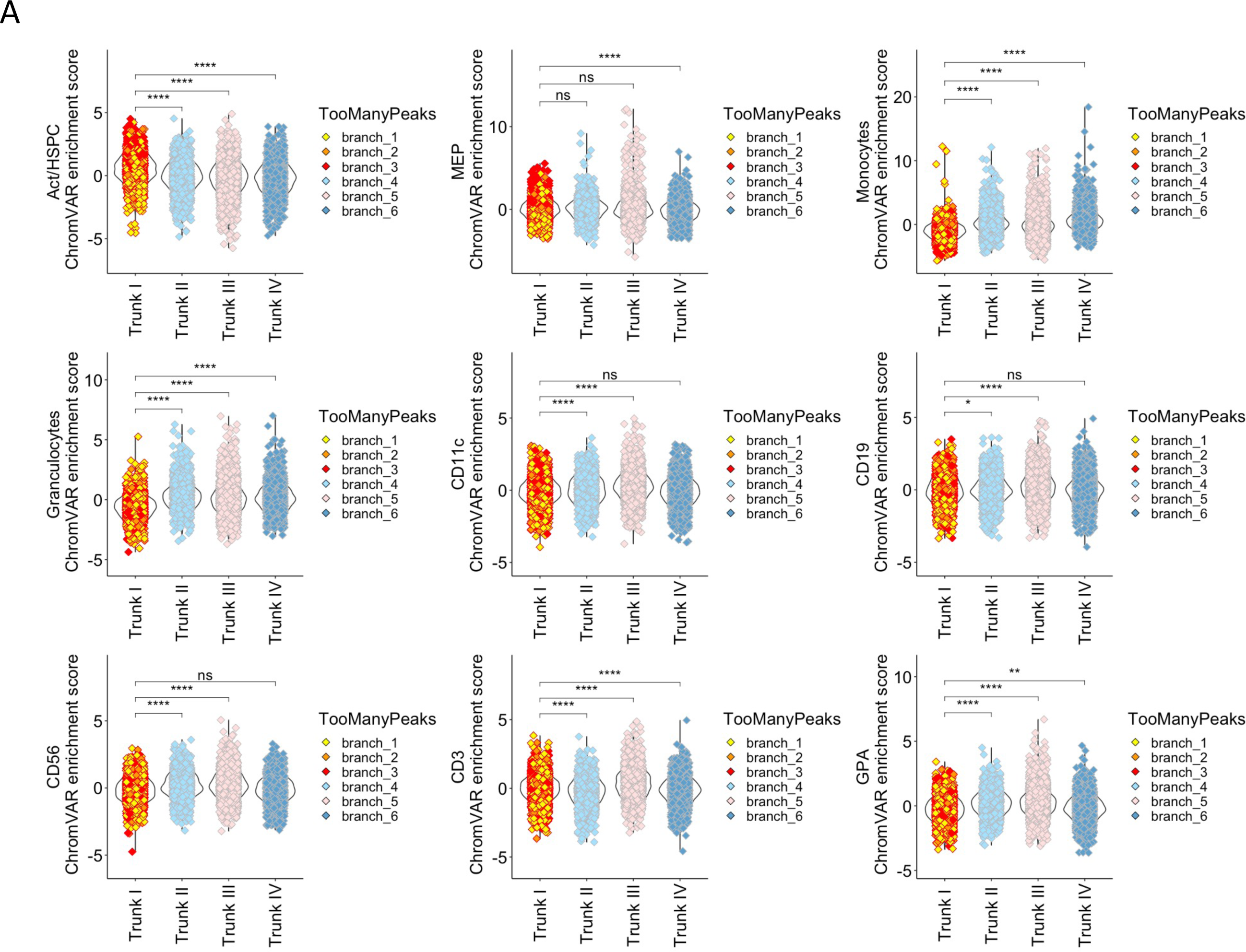
**A.** ChromVAR enrichment scores for each of the epigenetic signatures from ^42^ are disclosed for OCI-AML22 CD34+CD38- cells across the ToomanyCells branches as indicated. Cells are colored based on the Branch they were allocated to in the ToomanyPeaks Tree from Figure 1F.

**Supplementary Figure S6.**
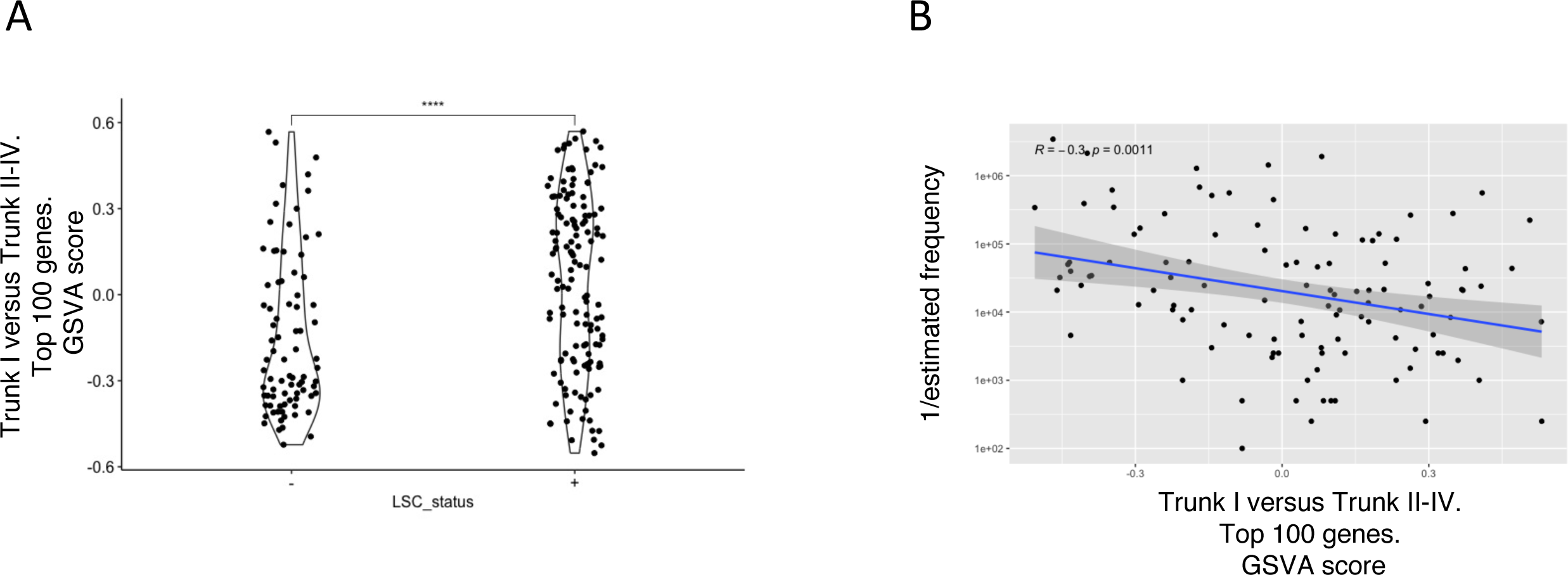
**A.** GSVA score was calculated on LSC depleted (LSC-) or LSC enriched (LSC+) fractions from primary AML samples ^2^ for the list of genes upregulated in TrunkI cells versus the other Trunks (Trunk II-IV) according to TooManyCells algorithm and described in Figure 1. **B.** For each LSC+ fractions extracted from primary AML samples^2^, GSVA score the list of genes upregulated in TrunkI cells versus the other Trunks (Trunk II-IV) according to TooManyCells algorithm and described in Figure 1 was calculated and plotted against the estimated LSC frequency assessed through in vivo assay^2^ (1/estimated frequency).

**Supplementary Figure S7.**
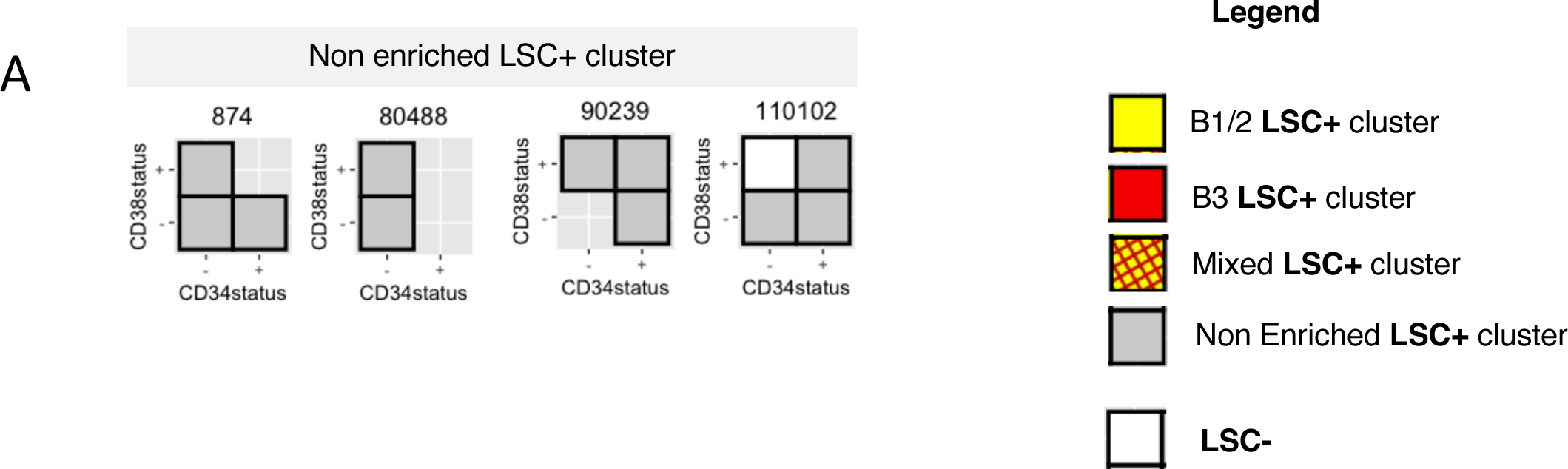
**A.** GSVA was performed for B1/2 or B3 signatures across functionally assessed LSC-positive fractions from primary AML samples^2^. Scores for each signature are plotted and were used to perform hierarchical clustering using Complexe heatmap. Functionally assessed LSC positive fractions for each indicated patient sample are represented in boxes colored based on the cluster each fraction was allocated in Figure 2A. LSC negative fractions are indicated in white. Not assessed fractions are not represented. Patients with fractions belonging to the non enriched LSC+ cluster from Figure 2A (A) are represented.

